# YAP/TAZ-controlled ERK dynamics coordinate progenitor expansion and differentiation commitment

**DOI:** 10.64898/2026.09.24.753950

**Authors:** Sanjeev Sharma, Henri Berger, Zhibo Zhang, Tobias Meyer, Mary N. Teruel

## Abstract

Progenitor cells must proliferate to expand the cell population, yet terminal differentiation requires this proliferative state to end. How signaling controls the duration of this proliferative window remains poorly understood. Using adipogenesis and live single-cell imaging of differentiation, cell-cycle, and ERK-activity reporters, we show that YAP and TAZ coordinate progenitor expansion with differentiation commitment by regulating ERK dynamics. YAP/TAZ maintain cells in a fluctuating high-ERK state that promotes proliferation while actively keeping the differentiation driver PPARG below the threshold for irreversible commitment. Crucially, this differentiation block is not explained by proliferation alone: inhibiting CDK4/6 or AKT suppressed proliferation without restoring differentiation, whereas MEK–ERK inhibition restored differentiation even when YAP/TAZ activity remained high. As YAP/TAZ activity decreases, dampened ERK fluctuations trigger PPARG activation. These findings support a self-limiting model in which YAP/TAZ-driven progenitor expansion progressively increases cell density and contact-dependent Hippo signaling, reducing YAP/TAZ activity and terminating the proliferative phase. Consequently, transient YAP/TAZ activation expands the progenitor pool while preserving subsequent differentiation, whereas sustained activation suppresses commitment. Together, these findings identify YAP/TAZ-controlled ERK dynamics as the nexus coordinating progenitor expansion with terminal differentiation and suggest that slower density-dependent Hippo feedback may set the duration of this proliferative window to regulate differentiated cell-number output.

## Introduction

Terminal differentiation requires progenitor cells to balance two opposing demands: they must proliferate enough to expand the progenitor pool, but they must eventually stop proliferating and commit to differentiation^1–5^. The number of differentiated cells produced therefore depends on how long this proliferative window remains open, because each additional division expands the pool of cells available to differentiate^2,4^. Yet the signaling mechanisms that determine the duration of this window remain poorly understood^3–5^.

Importantly, stopping proliferation is not sufficient to trigger differentiation. In adipocyte progenitors, we previously showed that commitment occurs when PPARG, the master regulator of adipogenesis, rises above a critical threshold^3^ and requires sequential inactivation of CDK4/6, CDK2, and finally ERK during G1^6^. Blocking CDK4/6 and CDK2 stops cell-cycle progression but is not sufficient to induce differentiation commitment^6^. Conversely, once PPARG rises above its commitment threshold, cells enter the terminal differentiation program rather than continuing through additional proliferative cycles^3^. Thus, controlling differentiated-cell number requires more than controlling cell-cycle exit. It requires a mechanism that sustains progenitor expansion for a limited period while delaying commitment, and then permits differentiation to proceed.

One candidate signaling regulator is YAP, a well-established promoter of progenitor proliferation that has also been linked to differentiation in several tissues. In the intestine, YAP activation expands undifferentiated progenitors, whereas loss of YAP activity permits differentiation to resume^7^. In pancreatic progenitors, sustained YAP activation increases proliferation and impairs endocrine differentiation, whereas YAP inhibition promotes endocrine and beta-cell differentiation^8^. These studies support a model in which high YAP activity suppresses differentiation. In adipogenesis, however, the relationship appears different. Yap overexpression in mice increased adipocyte formation and adipose tissue mass^9^. Thus, depending on the system, YAP activity has been associated with either inhibition or promotion of differentiation, leaving its role in controlling the transition from progenitor expansion to differentiation unresolved.

Adipogenesis provides a particularly useful system for addressing this problem. Adipocyte progenitors undergo a variable number of divisions before commitment, and both proliferation and differentiation can be followed in the same individual cells using live cell-cycle reporters and endogenous PPARG^3^. This makes it possible to determine how YAP activity controls the duration of progenitor expansion and how that control is linked to differentiation commitment. Adipogenesis is also especially relevant to YAP biology because Yap overexpression in mice caused a striking and relatively selective expansion of white adipose tissue, with much smaller effects on most other organs^9^.

Here, we manipulate the level and duration of YAP activity and test the contribution of its paralog TAZ while following proliferation and differentiation in individual cells. We show that YAP and TAZ together maintain adipocyte progenitors in a proliferative, low-PPARG state. Reducing YAP/TAZ activity accelerates differentiation, whereas transient YAP/TAZ activation expands the progenitor population and ultimately increases the number of differentiated cells produced. In contrast, sustained YAP/TAZ activation suppresses commitment. Mechanistically, YAP/TAZ do not suppress differentiation simply by driving cell-cycle progression. Instead, they maintain high, fluctuating ERK activity, which restrains PPARG accumulation. MEK–ERK inhibition removes this block and restores differentiation despite sustained YAP/TAZ activity. As YAP/TAZ activity declines, cells spend less time with high ERK activity, allowing PPARG to rise and commitment to proceed. Together, these findings identify YAP/TAZ-controlled ERK dynamics as a mechanism that sustains progenitor expansion while delaying differentiation commitment, thereby controlling the duration of the proliferative window and the number of differentiated cells ultimately produced.

## Results

### YAP and TAZ act together to promote progenitor expansion and delay differentiation commitment

To test the role of YAP and TAZ during terminal differentiation, we used adipogenesis as a model program. We previously generated OP9 preadipocytes in which endogenous PPARG, the master regulator of adipogenesis, is fluorescently tagged^3,10^, allowing PPARG dynamics to be followed live in individual cells. Adipogenic stimulation gradually increases PPARG through external signaling and positive-feedback mechanisms. Once PPARG rises above a critical threshold, it rapidly switches to a self-sustaining high state that marks irreversible differentiation commitment, even after the adipogenic stimulus is removed (Fig. 1a,b). Before reaching this threshold, progenitor cells can undergo a variable number of divisions. Once the threshold is crossed, cells permanently exit the cell cycle and enter the terminal adipocyte differentiation program. Thus, adipogenesis begins with a period of progenitor expansion that precedes differentiation commitment (Fig. 1c). Because each progenitor division doubles the number of cells available for subsequent differentiation, the number of divisions before commitment directly determines differentiated-cell output.

**Figure 1.**
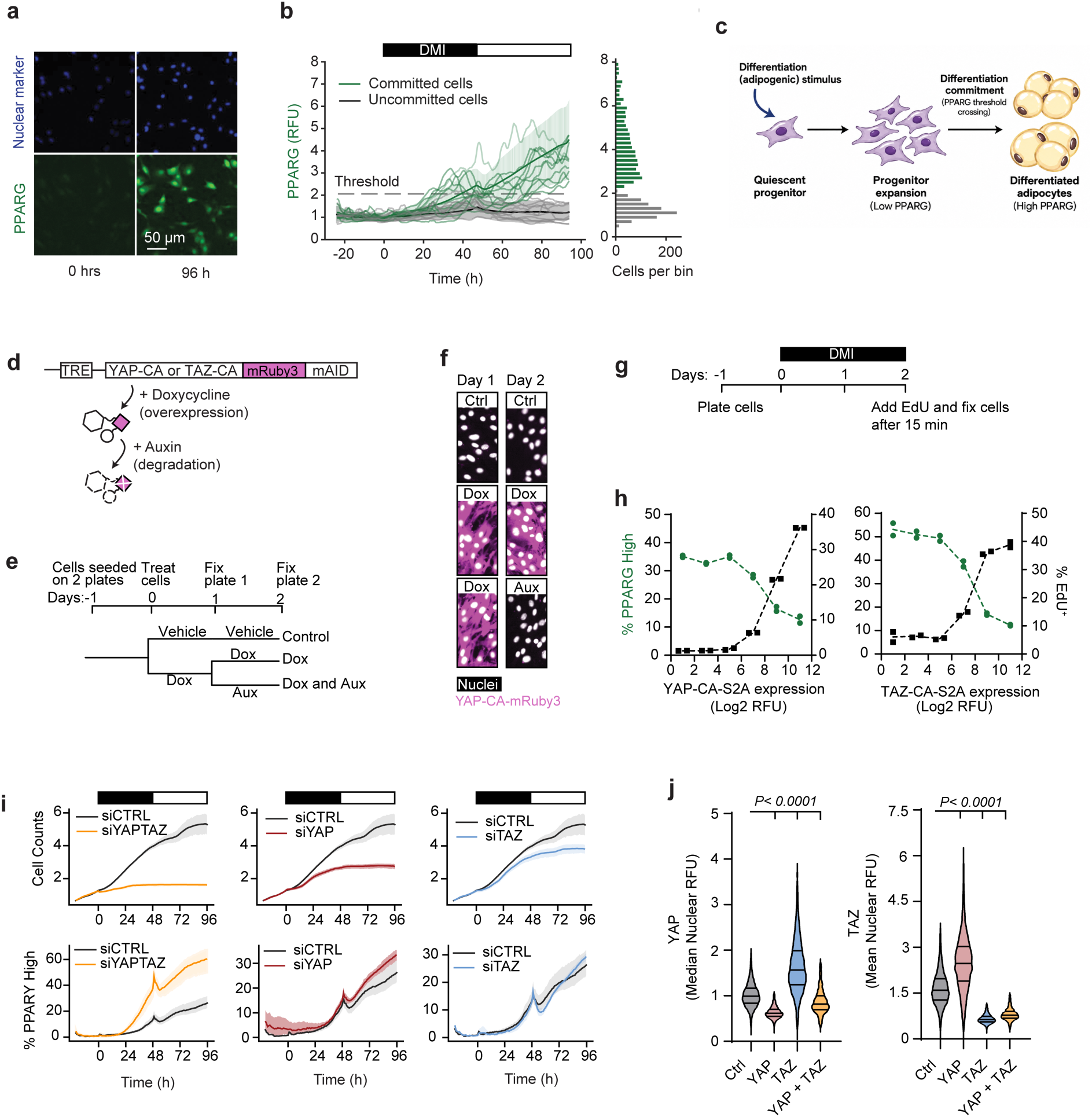
YAP and TAZ act together to promote progenitor expansion and delay differentiation commitment. (a) Immunofluorescence images of OP9 cells expressing endogenously tagged PPARG-mCitrine (green) and a nuclear marker (blue) before (0 h) and after 96 h of adipogenic differentiation. Scale bar, 50 µm. (b) Single-cell PPARG-mCitrine trajectories during adipogenic differentiation. Cells that crossed the PPARG commitment threshold are shown in green and uncommitted cells in gray. The dashed line indicates the commitment threshold. DMI was present for the first 48 h (bar). Right, distribution of PPARG levels at 96 h. (c) Schematic illustrating progenitor expansion before adipogenic differentiation commitment. Following an adipogenic stimulus, low-PPARG progenitors can undergo multiple divisions, expanding the population available for subsequent differentiation. Commitment occurs when PPARG crosses its threshold, after which cells proceed to the PPARG-high differentiated adipocyte state. (d) Design of the doxycycline-inducible, auxin-degradable constitutively active YAP and TAZ constructs (YAP-CA or TAZ-CA–mRuby3–mAID). Doxycycline induces construct expression, whereas auxin promotes degradation of the induced protein. (e) Experimental scheme used to validate reversible YAP-CA expression. Cells were treated with vehicle or doxycycline beginning on day 0. Expression was assessed on day 1. From day 1 to day 2, cells were maintained with vehicle or doxycycline, or auxin was added to doxycycline-treated cells while doxycycline was maintained. (f) Representative images of YAP-CA-mRuby3 (magenta) and nuclei (white) on days 1 and 2 for the indicated treatment conditions, showing low basal expression in control cells, induction by doxycycline, maintenance of expression with continued doxycycline, and loss of induced protein after auxin addition in the continued presence of doxycycline. (g) Experimental timeline for measuring proliferation and differentiation after YAP-CA or TAZ-CA induction. Cells were stimulated with DMI for 48 h. EdU was added for 15 min immediately before fixation on day 2. (h) Cells were treated with increasing doxycycline concentrations to generate a range of YAP-CA-S2A (left) or TAZ-CA-S2A (right) expression levels and binned according to nuclear mRuby3 fluorescence. Points show the mean percentage of PPARG-high cells (green, left axis) and EdU-positive cells (black, right axis) across two replicate wells for each expression bin. Data are representative of two independent experiments. (i) Total cell number (top) and percentage of PPARG-high cells (bottom) during 96 h of adipogenic differentiation after knockdown of YAP and TAZ together (siYAPTAZ), YAP alone (siYAP), or TAZ alone (siTAZ), compared with control siRNA (siCTRL). DMI was present for the first 48 h (bar). Line plots show mean ± 95% CI from three replicate wells per condition and are representative of two independent experiments. (j) Violin plots showing the distribution of single-cell nuclear YAP (left) and TAZ (right) fluorescence after the indicated siRNA treatments. Lines indicate the 25th, 50th, and 75th percentiles. At least 1,500 cells were analyzed per condition. Statistical significance was assessed by one-way ANOVA with Fisher’s LSD post-hoc test. All pairwise comparisons were performed against Ctrl siRNA condition, *P* < 0.0001.

To manipulate YAP/TAZ activity acutely, we generated doxycycline-inducible, auxin-degradable constructs expressing constitutively active YAP (YAP-CA-S2A) or TAZ (TAZ-CA-S2A)^11–13^ (Fig. 1d). We first validated that this system could reversibly control YAP/TAZ abundance (Fig. 1e,f). Doxycycline induced robust expression within 24 hours, whereas subsequent auxin addition rapidly depleted the induced protein despite continued doxycycline treatment. We then used this system to determine how increasing YAP or TAZ activity affects proliferation and differentiation. Cells were stimulated with DMI and exposed to increasing doxycycline concentrations to generate a range of YAP-CA or TAZ-CA expression levels, while proliferation was measured by EdU incorporation and differentiation by PPARG expression (Fig. 1g,h). Increasing either YAP or TAZ produced opposing changes in cell fate: the fraction of EdU-positive cells increased as the fraction of PPARG-high cells decreased (Fig. 1h). Thus, increasing either YAP or TAZ activity is sufficient to maintain progenitor proliferation while delaying differentiation commitment.

Because YAP and TAZ are closely related paralogs, we asked whether they act redundantly in controlling progenitor proliferation and differentiation commitment. Simultaneous knockdown of YAP and TAZ sharply reduced proliferation and nearly doubled the fraction of differentiated cells, whereas knockdown of either paralog alone had only modest effects. YAP knockdown increased TAZ abundance, whereas TAZ knockdown increased YAP abundance (Fig. 1j), suggesting reciprocal compensation between the two paralogs. Consistent with reciprocal regulation, constitutively active YAP reduced TAZ abundance, whereas constitutively active TAZ reduced YAP abundance (Supplementary Fig. 1). Together, these results suggest that YAP and TAZ buffer one another, helping explain why simultaneous loss of both is required to strongly shift cells from progenitor proliferation toward differentiation.

### YAP/TAZ–TEAD activity decreases as cells progress toward adipogenic commitment

Because increasing YAP/TAZ activity delays differentiation commitment, we next asked whether endogenous YAP/TAZ activity normally declines as cells approach commitment. Yap1 and Wwtr1 (TAZ) mRNA did not decrease across the four-day time course (Fig. 2a), indicating that reduced pathway output is not explained by transcriptional loss of either effector. We therefore measured YAP and TAZ protein localization by quantitative immunofluorescence. Total YAP decreased modestly and redistributed from nucleus to cytosol (Fig. 2b), causing the YAP nuclear-to-cytosolic ratio to fall from approximately 4.8 at day 0 to 1.8 at day 4 (Fig. 2c). TAZ showed a different change in abundance: total TAZ increased during differentiation, driven largely by accumulation in the cytosol, while nuclear TAZ remained approximately constant (Fig. 2e). Its nuclear-to-cytosolic ratio nevertheless fell over a similar range, from approximately 3.5 to 1.6 (Fig. 2f). Consistent with reduced nuclear retention, inhibitory YAP Ser127 phosphorylation increased after DMI stimulation^11^.

**Figure 2.**
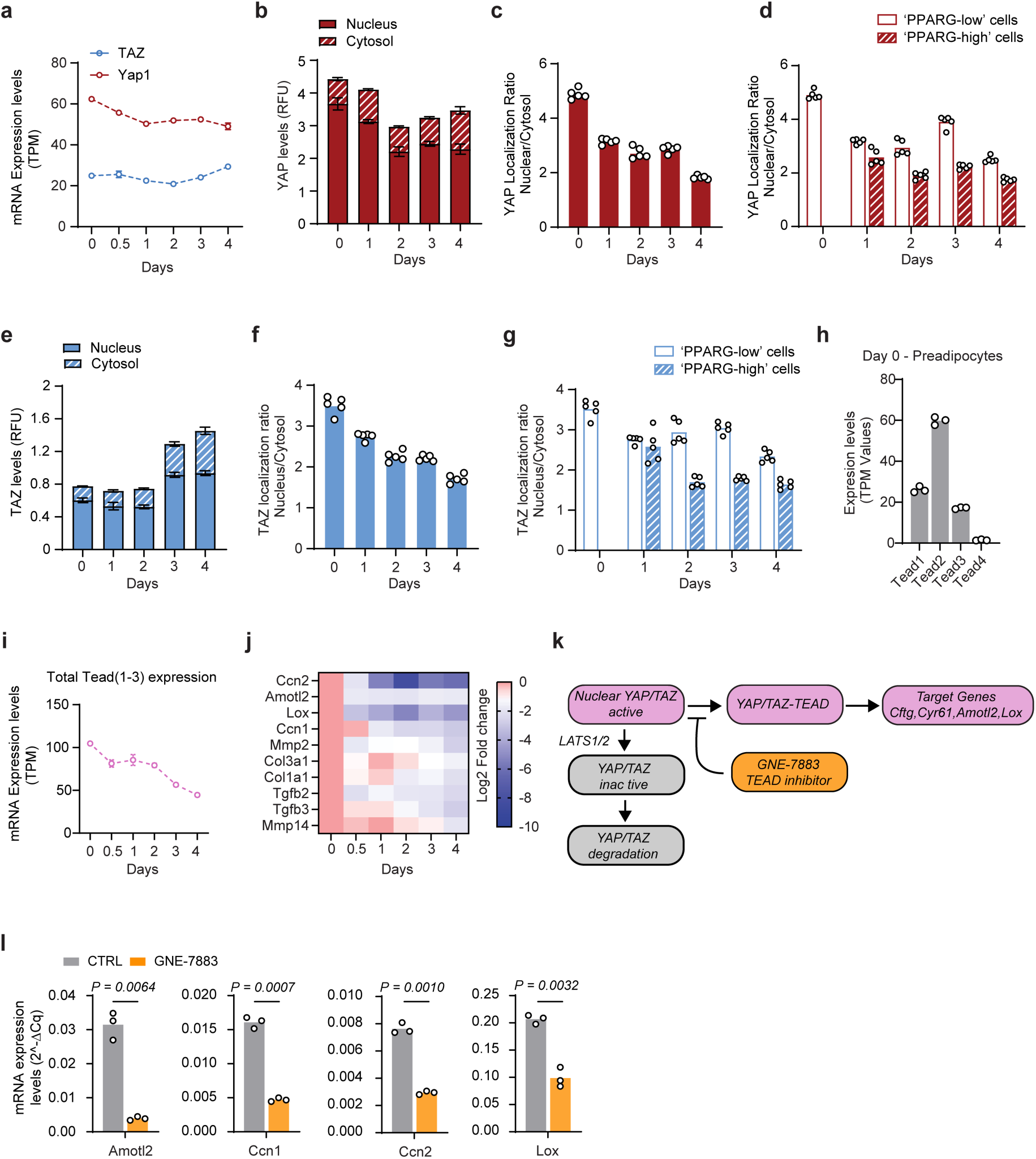
YAP/TAZ–TEAD activity decreases as cells progress toward adipogenic commitment. Unless otherwise indicated, cells were differentiated with DMI from day 0 to day 2, followed by insulin-containing medium without DMI from day 2 to day 4. (a) Yap1 and Wwtr1 (TAZ) mRNA levels (transcripts per million, TPM) measured by RNA-seq in OP9 cells over days 0–4 of adipogenic differentiation. Data are mean ± s.e.m. from three biologically independent samples per time point. (b) Total YAP protein (RFU) measured in single cells as the sum of nuclear (solid) and cytosolic (hatched) fractions by quantitative immunofluorescence over days 0–4 of adipogenesis. Bar plots show mean ± s.d. from 5 replicate wells; representative of two biologically independent experiments, ≥1,000 cells per experiment. (c) Mean nuclear-to-cytosolic ratio of YAP protein in single cells for data in Fig. 2b. Bars show the overall means for each time point, and individual points show the means from replicate wells. (d) Mean YAP nuclear-to-cytosolic ratio from PPARG-high and PPARG-low cells at each time point. Day 0 contains no PPARG-high cells for data shown in Fig. 2b. PPARG-high cells have significantly lower ratios than PPARG-low cells at all time points tested; Plotted as in Fig. 2c. Multiple t-tests (Day 1, *P* = 0.0022; Day 2-4, *P* < 0.0001). (e) Total TAZ protein (RFU) measured in single cells as the sum of nuclear (solid) and cytosolic (hatched) fractions by quantitative immunofluorescence over days 0–4 of adipogenesis. Total TAZ increases over the time course, driven by cytosolic accumulation, while nuclear TAZ remains approximately unchanged. Bars show mean ± s.d. from 5 replicate wells; representative of two biologically independent experiments, ≥1,000 cells per experiment. (f) Mean TAZ nuclear-to-cytosolic ratio in single cells over days 0–4 for data shown in Fig. 2e; plotted as in Fig. 2c. (g) TAZ nuclear-to-cytosolic ratio separated by PPARG level for data shown in Fig. 2e; plotted as in Fig. 2d. (h) *Tead1–Tead4* mRNA (TPM) measured by RNAseq analysis in unstimulated OP9 preadipocytes. Bars show the mean and points show the individual TPM values measured for three independent biological samples. (i) Combined *Tead1–3* mRNA levels (TPM) over days 0–4 of adipogenic differentiation of OP9 cells. Plots show the mean ± s.e.m. of three biologically independent samples per time point. (j) Heatmap of log2 fold change relative to day 0 for canonical YAP/TAZ–TEAD target genes (*Ccn2(Ctgf), Amotl2*, *Lox*, *Ccn1 (Cyr61)*) and extracellular-matrix genes in OP9 cells across days 0–4 of adipogenic differentiation measured by RNAseq. n = 3 biologically independent samples per time point. (k) Scheme showing two levels of YAP/TAZ–TEAD regulation: LATS1/2-dependent cytoplasmic redistribution reduces the active nuclear pool of YAP and TAZ, and declining TEAD expression limits transcriptional output; GNE-7883 blocks YAP/TAZ–TEAD-dependent transcription. (l) Expression of the canonical YAP/TAZ–TEAD target genes *Amotl2*, *Ccn1 (Cyr61)*, *Ccn2 (Ctgf)*, and *Lox* after 24 h of GNE-7883 treatment (2.5 µM) or vehicle (DMSO), measured by qPCR and normalized to *Gapdh*. *n* = 3 independently treated biological samples, each measured in technical duplicate. Technical duplicates were averaged within each biological sample before statistical analysis. Bars show the mean and points indicate individual biological-sample 2^-ΔCq values. Unpaired two-tailed *t*-tests with Welch’s correction; *Amotl2*, *P* = 0.0064; *Ccn1*, *P* = 0.0007; *Ccn2*, *P* = 0.0010; *Lox*, *P* = 0.0032.

Because differentiation is asynchronous, population averages combine cells at different stages of the program. We therefore separated cells at each time point according to single-cell PPARG level. At matched time points, PPARG-high cells consistently had lower YAP and TAZ nuclear-to-cytosolic ratios than PPARG-low cells (Fig. 2d,g). The lower nuclear-to-cytosolic ratios in PPARG-high cells link the decline in YAP/TAZ nuclear localization to differentiation state rather than simply to time after adipogenic stimulation. The perturbation experiments below test whether this relationship is functional.

We next asked whether reduced TEAD availability provides an additional mechanism for lowering YAP/TAZ transcriptional output. Tead1–3 were the TEAD paralogs expressed in day-0 preadipocytes, whereas Tead4 was essentially absent (Fig. 2H). Combined Tead1–3 transcript levels then declined progressively during differentiation (Fig. 2I). In parallel, canonical YAP/TAZ–TEAD target genes Ccn2 (Ctgf), Ccn1 (Cyr61), Amotl2, and Lox decreased across the same time course, together with several extracellular-matrix genes^14,15^ (Fig. 2j).

A scheme summarizes these two levels of regulation: changes in YAP/TAZ localization reduce the active nuclear pool, while declining TEAD expression limits YAP/TAZ-dependent transcription (Fig. 2k). Consistent with the latter, acute inhibition of TEAD with GNE-7883 reduced Ccn2, Ccn1, Amotl2, and Lox expression^14,16^ (Fig. 2l). Together, these results show that the YAP/TAZ–TEAD transcriptional program is progressively dismantled as cells approach commitment, through both loss of nuclear YAP/TAZ and reduced TEAD availability. This progressive decline provides a mechanism by which the YAP/TAZ-dependent proliferative state can be terminated as cells approach commitment.

### YAP/TAZ activity controls progenitor expansion and differentiated-cell output

We next asked whether changing endogenous YAP/TAZ activity alters how many divisions cells undergo before differentiating. YAP/TAZ activity was reduced by knocking down YAP and TAZ and increased by knocking down LATS1, and proliferation and differentiation were followed by live imaging for four days (Fig. 3a). YAP/TAZ knockdown accelerated differentiation, whereas LATS1 knockdown delayed it (Fig. 3b). At the end of the experiment, YAP/TAZ knockdown produced fewer total PPARG-high cells, whereas LATS1 knockdown produced more (Fig. 3c). Division counting showed why: YAP/TAZ knockdown shifted cells toward fewer divisions, whereas LATS1 knockdown increased the number of cells undergoing multiple divisions before commitment (Fig. 3d-f). Thus, endogenous YAP/TAZ activity does not simply alter the fraction of cells that differentiate. It controls differentiated-cell output by determining how many progenitor divisions occur before commitment.

**Figure 3.**
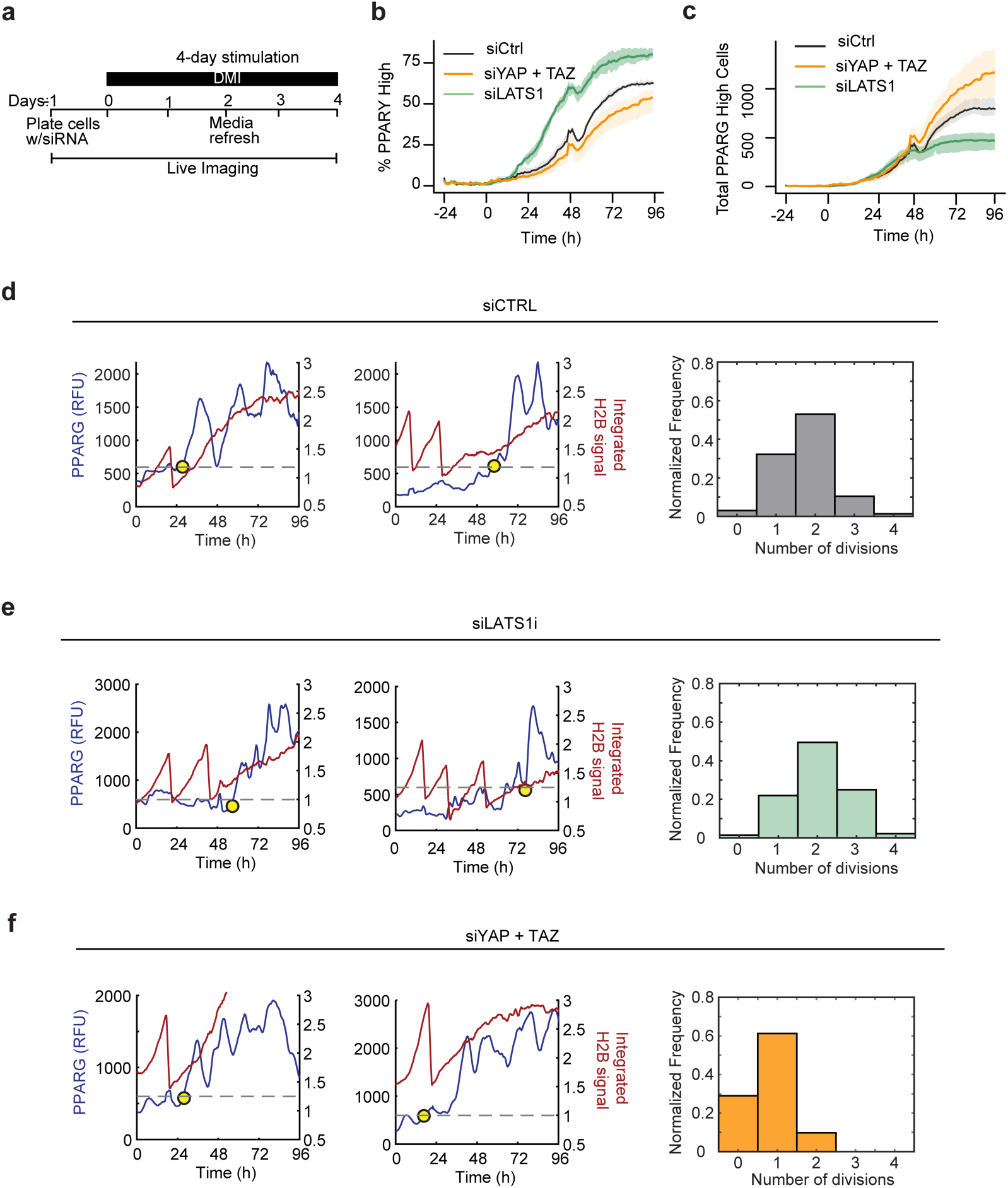
Endogenous YAP/TAZ activity controls progenitor expansion and differentiated-cell output. (a) Experimental timeline. OP9 cells were transfected with siRNA and plated on day −1. Adipogenic differentiation was initiated with DMI on day 0, and cells were imaged continuously for 96 h, with a media refresh on day 2. DMI was maintained throughout the 96-h experiment. (b) Percentage of PPARG-high cells over time after treatment with control siRNA (siCTRL), combined YAP and TAZ siRNAs (siYAP + TAZ), or LATS1 siRNA (siLATS1). Line plots show mean ± s.e.m. from six replicate wells per condition. (c) Total number of PPARG-high cells over time for the conditions shown in b. Line plots show mean ± s.e.m. from six replicate wells per condition. (d–f) Single-cell analysis of PPARG accumulation, cell division, and differentiation commitment after treatment with siCTRL (d), siLATS1 (e), or siYAP + TAZ (f). Left and middle panels show representative single-cell traces of PPARG (blue, left axis) and integrated H2B signal (red, right axis). Abrupt decreases in integrated H2B signal mark cell divisions, and yellow circles indicate the time of differentiation commitment. Right, normalized frequency distributions of the number of divisions completed before differentiation commitment.

### A transient, but not continuous, pulse of YAP/TAZ activity increases differentiated-cell output

The siRNA experiments changed YAP/TAZ activity throughout differentiation, but they could not distinguish whether YAP/TAZ activity must simply be low for differentiation or instead must be high early and then fall. We therefore tested whether a transient early increase in YAP/TAZ could expand the progenitor pool without blocking later commitment. Using the inducible-degradable constructs, we compared continuous activation across the four-day program with a one-day pulse initiated with DMI and terminated by auxin-mediated degradation (Fig. 4a). We tested YAP-CA-S2A, the more strongly active YAP-CA-S5A, and TAZ-CA-S2A (Fig. 4b–m).

**Figure 4.**
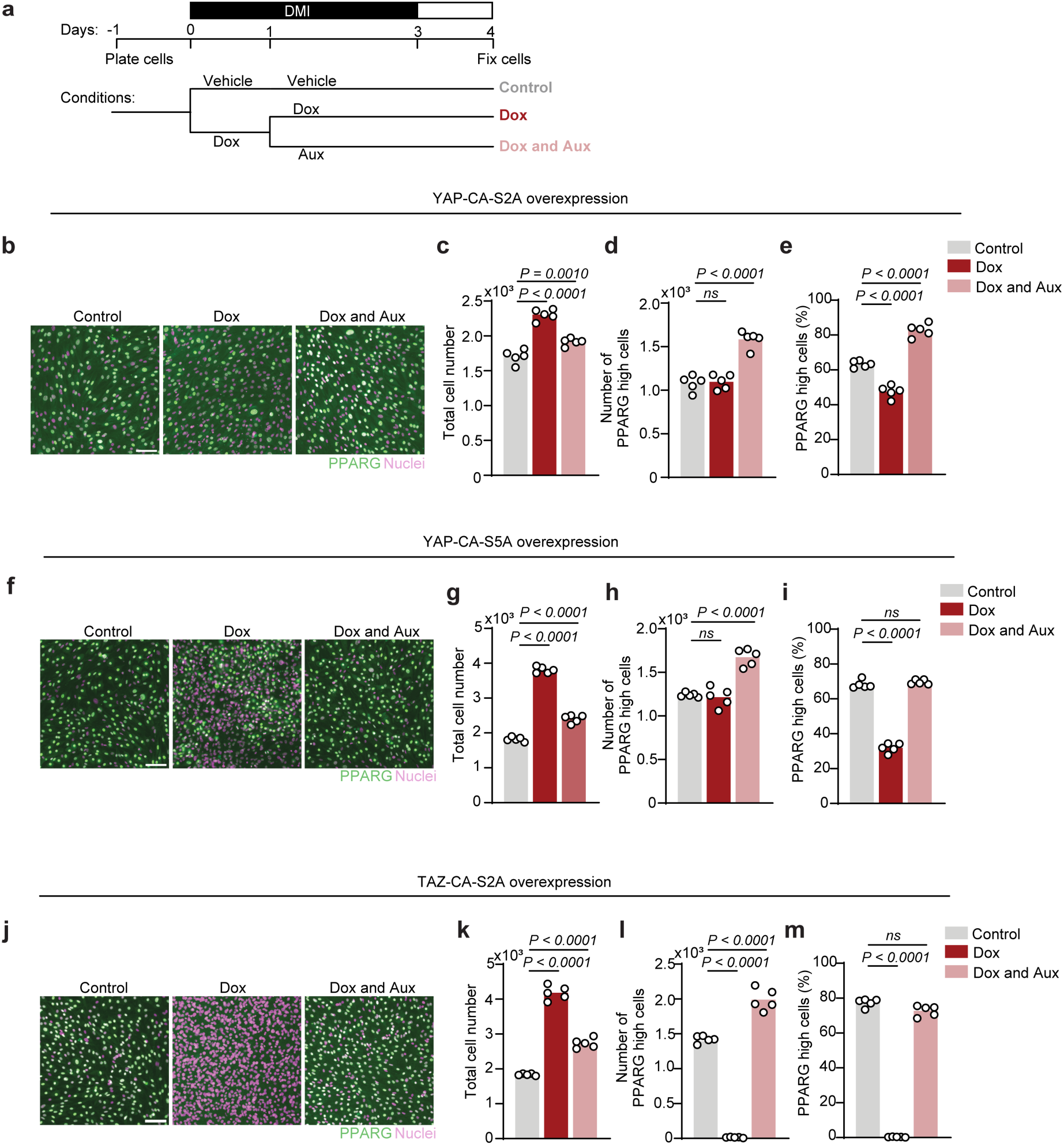
A transient pulse of YAP/TAZ activation expands progenitors without blocking terminal differentiation. **(a)** Experimental timeline and treatment conditions. OP9 cells expressing doxycycline-inducible, auxin-degradable YAP-CA or TAZ-CA constructs were plated on day −1 and stimulated with DMI from day 0 to day 3. Cells were treated with vehicle throughout (Control), doxycycline from day 0 to day 4 (Dox), or doxycycline from day 0 to day 1, after which doxycycline was removed and auxin was added from day 1 to day 4 (Dox and Aux). Cells were fixed on day 4. Doxycycline concentrations were 3 µg/mL for YAP-CA-S2A, 1 µg/mL for YAP-CA-S5A, and 0.25 µg/mL for TAZ-CA-S2A. **(b)**, **(f)**, **(j)** Representative images of PPARG (green) and nuclei (magenta) for Control, Dox, and Dox + Aux conditions for YAP-CA-S2A (**b**), YAP-CA-S5A (**f**), and TAZ-CA-S2A (**j**). Scale bar, 100 µm. **(c)**, **(g)**, **(k)** Total cell number for YAP-CA-S2A (**c**), YAP-CA-S5A (**g**), and TAZ-CA-S2A (**k**). **(d)**, **(h)**, **(l)** Number of PPARG-high cells for YAP-CA-S2A (**d**), YAP-CA-S5A (**h**), and TAZ-CA-S2A (**l**). **(e)**, **(i)**, **(m)** Percentage of PPARG-high cells for YAP-CA-S2A (e), YAP-CA-S5A (i), and TAZ-CA-S2A (m). For c–e, g–i, and k–m, bars show the mean and points indicate individual replicate wells, with five replicate wells per condition. Data are representative of two independent experiments. Statistical significance was assessed by one-way ANOVA followed by Dunnett’s multiple-comparisons test. Exact P values are shown in the figure; ns, not significant.

Continuous YAP or TAZ activation increased total cell number but reduced the fraction of cells that differentiated (Fig. 4c,e,g,i,k,m). For the YAP constructs, these opposing effects largely offset one another, producing a similar total number of differentiated cells as control. Continuous TAZ activation suppressed differentiation more strongly, consistent with additional TAZ-mediated repression of PPARG^17^. In contrast, a transient pulse of YAP or TAZ increased total cell number while preserving or increasing the differentiated fraction, producing more PPARG-high cells from the same starting population (Fig. 4d,h,l). A transient pulse therefore allows additional progenitor divisions without preventing the later rise in PPARG and commitment. This demonstrates that the effect of YAP/TAZ depends on duration: transient activity can increase final adipocyte output by expanding the progenitor pool, whereas sustained activity continues to delay the transition to differentiation.

### YAP/TAZ suppress differentiation through ERK rather than proliferation alone

To determine how YAP/TAZ suppress differentiation, we used the LATS1/2 inhibitor TDI-011536, which increases nuclear YAP/TAZ abundance^18^ (Fig. 5a,b). LATS inhibition increased cell number and reduced differentiation, and both effects were reversed by TEAD inhibition (Fig. 5C). Because TEAD inhibition reversed both effects, LATS inhibition acts through YAP/TAZ–TEAD to increase proliferation and suppress differentiation.

**Figure 5.**
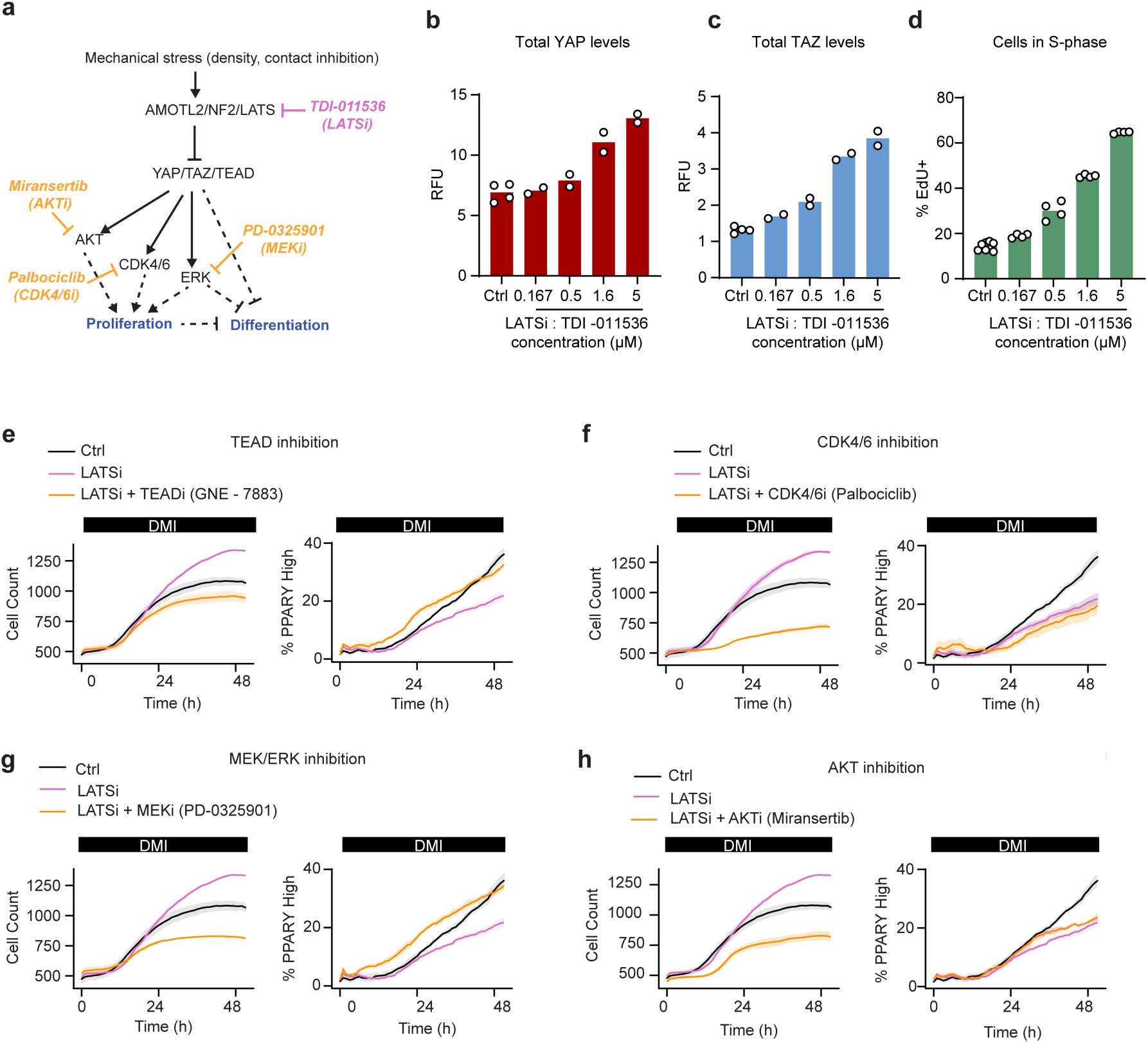
YAP/TAZ suppress differentiation through MEK–ERK rather than through proliferation alone. (a) Working model and perturbation strategy used to distinguish the proliferative and differentiation-suppressive effects of YAP/TAZ. Mechanical inputs, including cell density and contact inhibition, act through AMOTL2/NF2/LATS to restrain YAP/TAZ–TEAD activity. YAP/TAZ–TEAD signaling promotes proliferation through AKT and CDK4/6 and suppresses differentiation through ERK. Pharmacological inhibitors used to perturb these pathways are indicated in color. **(b), (c)** Total YAP (in **b**) and TAZ (in **c**) levels (nuclear + cytosol) measured by quantitative immunofluorescence after 24 h treatment with TDI-011536 at the indicated concentrations versus untreated control. Bars show the overall means and data points show the individual means of 2-4 replicate wells per condition. n ≥ 1,000 cells per condition. **(d)** Percentage of EdU-positive cells after 24 h treatment with TDI-011536 at the indicated concentrations. Bars show the mean, and points indicate individual replicate-well means. **(e)** – **(h)** Dual-reporter OP9 cells expressing PPARG-mCitrine and an mCherry-APC/C cell-cycle reporter were stimulated with DMI and treated with vehicle (Ctrl), the LATS1/2 inhibitor TDI-011536 (LATSi), or LATSi together with the indicated pathway inhibitor: TEAD inhibitor GNE-7883 (e), CDK4/6 inhibitor palbociclib (f), MEK inhibitor PD-0325901 (g), or AKT inhibitor miransertib (h). Cells were imaged for 48 h. Left, total cell number. Right, percentage of PPARG-high cells. Lines show the mean and shaded regions indicate 95% confidence intervals across three replicate wells per condition. The Ctrl and LATSi traces are the same dataset in e–h and are repeated to facilitate comparison with each inhibitor condition. Inhibitor concentrations were 200 nM TDI-011536, 2.5 µM GNE-7883, 2.5 µM palbociclib, 200 nM PD-0325901, and 200 nM miransertib.

Because YAP induces Cyclin D and CDK4 activity^19^, we first asked whether YAP/TAZ block differentiation simply by driving proliferation. Palbociclib strongly reduced the proliferation induced by LATS inhibition but did not restore differentiation (Fig. 5d). In contrast, MEK inhibition suppressed the increase in cell number and restored differentiation (Fig. 5E). This result is consistent with our previous finding that ERK suppresses PPARG commitment even after CDK4/6 and CDK2 are inhibited^6^. MEK–ERK effects on adipogenesis are stage dependent: early ERK activity can promote adipogenic gene expression^20^, whereas ERK-dependent phosphorylation can inhibit PPARG activity^21,22^.

Because CDK4/6 and CDK2 act redundantly in these cells^6^, we used AKT inhibition as a second independent way to suppress the YAP/TAZ-driven increase in proliferation. AKT inhibition reduced proliferation to a similar extent as MEK inhibition but did not restore differentiation (Fig. 5F–H). Constitutively active YAP or TAZ also increased phospho-ERK without increasing total ERK (Supplementary Fig. 3). CDK4/6 and AKT inhibition therefore reduced proliferation without restoring differentiation, whereas MEK inhibition restored differentiation despite a similar reduction in cell number. YAP/TAZ therefore suppress PPARG commitment through ERK rather than through proliferation alone. These experiments separate the proliferative and differentiation-suppressive functions of YAP/TAZ and identify ERK as the signaling branch that links YAP/TAZ activity to differentiation commitment.

### YAP/TAZ shift ERK dynamics toward a high-activity state to delay commitment

The inhibitor experiments identified ERK as the signaling branch through which YAP/TAZ suppress commitment, but they did not reveal how ERK activity changes in individual cells during this transition. We therefore measured ERK dynamics directly during differentiation by combining the PPARG reporter with an ERK kinase-translocation reporter (ERK-KTR), whose cytoplasmic-to-nuclear ratio reports ERK activity (38) (Fig. 6a,b). Single-cell traces showed rapid ERK fluctuations and substantial cell-to-cell variability. During adipogenic stimulation, cells progressively shifted from spending more time at high ERK activity toward a lower-ERK state before PPARG rose (Fig. 6c,d). Rather than switching abruptly from an ERK-on to an ERK-off state, cells progressively reduced the fraction of time spent at high ERK activity before commitment.

**Figure 6.**
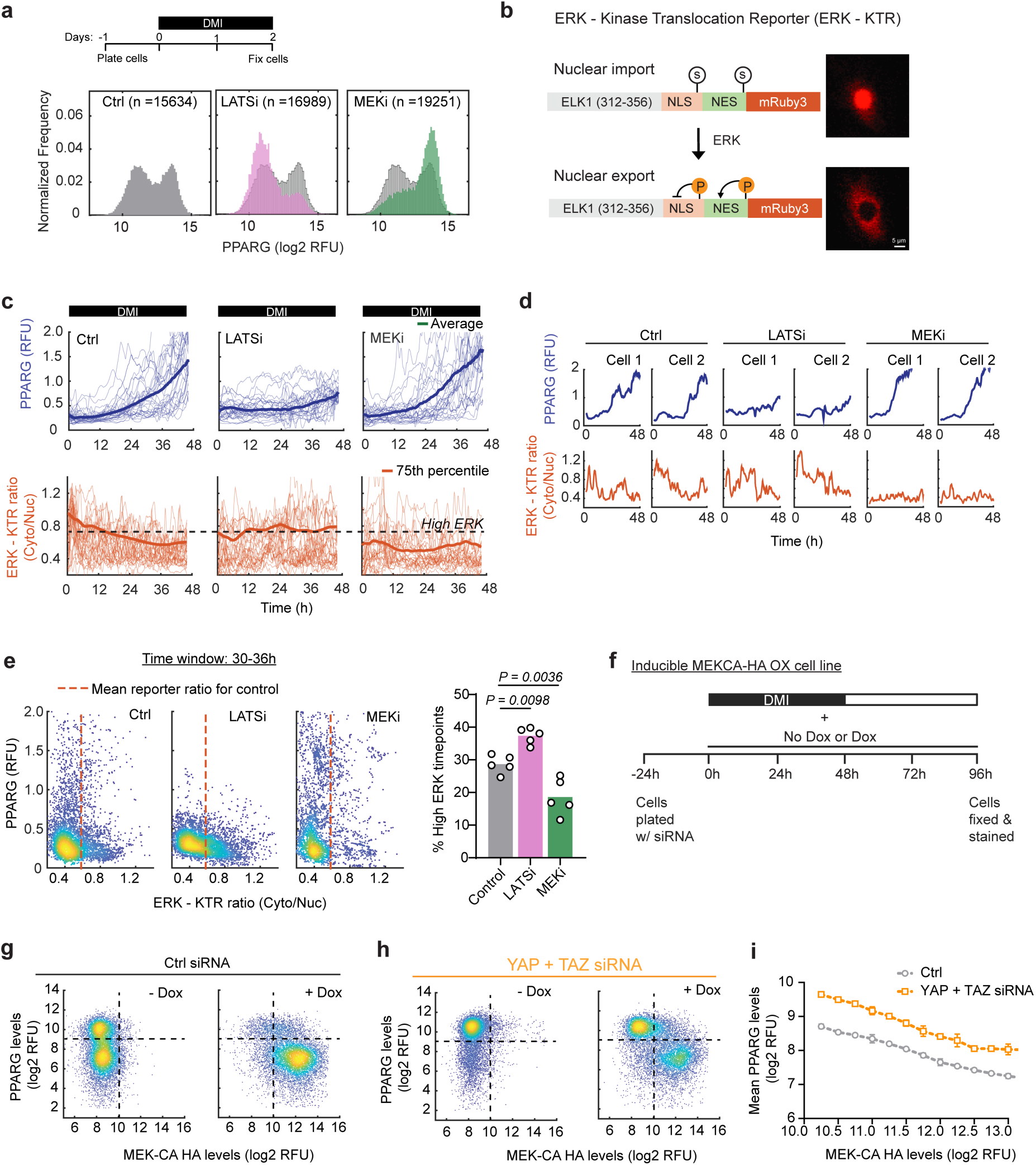
YAP/TAZ increase the fraction of time cells spend in the high-ERK state to delay commitment. (a) Experimental timeline. Cells were stimulated with DMI in the presence of vehicle, the LATS1/2 inhibitor TDI-011536 (LATSi), or the MEK inhibitor PD-0325901 (MEKi) and imaged live for 48 h. Histograms show the distribution of PPARG levels at 48 h for each condition. (b) Design of the ERK kinase-translocation reporter (ERK-KTR). The reporter contains an ERK-responsive ELK1 substrate fused to nuclear localization and export sequences and mRuby3. ERK-dependent phosphorylation promotes nuclear export, such that the cytoplasmic-to-nuclear fluorescence ratio reports ERK activity. Representative images illustrate predominantly nuclear localization at low ERK activity and cytoplasmic localization at high ERK activity. Scale bar, 5 µm. (c) Population dynamics of PPARG (top) and ERK-KTR activity (bottom) during 48 h of adipogenic differentiation in control, LATSi (TDI-011536, 200 nM), and MEKi (PD-0325901, 200 nM) conditions. Thin lines show individual-cell trajectories. For PPARG, the bold line indicates the population average. For ERK-KTR, the bold line indicates the 75th percentile of the single-cell distribution. The dashed horizontal line indicates the threshold used to define the high-ERK state. (d) Representative single-cell PPARG (blue, top) and ERK-KTR (orange, bottom) trajectories from two cells per condition. PPARG accumulation occurs after cells transition away from sustained high ERK activity. (e) Left, single-cell PPARG levels plotted against ERK-KTR activity during the 30–36 h time window for control, LATSi, and MEKi conditions. The vertical dashed line indicates the ERK-KTR threshold used to define high ERK activity. Right, percentage of time points classified as high ERK during the same time window. Bars show the mean, and points indicate individual replicate wells (n = 5 wells per condition). Statistical significance was assessed by one-way ANOVA followed by Dunnett’s multiple-comparisons test, with each treatment compared with control. Exact P values are shown. (f) Experimental timeline for testing whether constitutive MEK activation can bypass loss of YAP/TAZ. OP9 PPARG-mCitrine cells expressing doxycycline-inducible HA-tagged constitutively active MEK1 (MEK-CA-HA) were transfected with control or YAP + TAZ siRNA 24 h before DMI stimulation. DMI was present for the first 48 h. Cells were treated with increasing doxycycline concentrations to generate a range of MEK-CA-HA expression levels and were fixed at 96 h for HA immunofluorescence and PPARG-mCitrine measurement. (g, h) Single-cell PPARG-mCitrine levels plotted against MEK-CA-HA expression at 96 h in control siRNA (g) and YAP + TAZ siRNA (h) conditions, with and without doxycycline. Horizontal dashed lines indicate the PPARG-high threshold, and vertical dashed lines indicate the threshold for induced MEK-CA-HA expression. (i) Cells from the doxycycline-treated conditions were binned according to MEK-CA-HA expression (0.5 log2 RFU bins), and mean PPARG levels were calculated for each bin. Points show the mean PPARG level across three replicate wells, with error bars indicating s.d., for control and YAP + TAZ siRNA conditions.

Increasing YAP/TAZ activity by LATS inhibition kept cells in the high-ERK state for longer, whereas MEK inhibition maintained low ERK activity (Fig. 6c–e). Across conditions, the fraction of PPARG-high cells varied inversely with the amount of time cells spent at high ERK activity in the 30–36-h window (Fig. 6e). Because ERK activity declined before PPARG rose and LATS inhibition prolonged the time spent at high ERK activity, these data support a model in which YAP/TAZ delay commitment by keeping cells at high ERK activity for longer.

If YAP/TAZ act upstream of MEK–ERK to suppress commitment, then constitutively activating MEK should bypass the loss of YAP/TAZ and restore suppression of PPARG. We tested this prediction using a doxycycline-inducible constitutively active MEK1 construct (MEK-CA-HA) in control and YAP + TAZ siRNA-treated cells (Fig. 6f). YAP/TAZ knockdown increased the fraction of PPARG-high cells in the absence of MEK-CA induction (Fig. 6g,h). Constitutively active MEK suppressed PPARG in both control and YAP/TAZ-depleted cells, and PPARG levels decreased as MEK-CA expression increased (Fig. 6g–i). Thus, activated MEK bypasses the requirement for YAP/TAZ, placing YAP/TAZ function upstream of MEK–ERK in the pathway controlling differentiation commitment.

Together, these results place MEK–ERK downstream of YAP/TAZ and show that the amount of time cells spend at high ERK activity links YAP/TAZ activity to differentiation commitment. High YAP/TAZ promotes progenitor expansion and prolongs high ERK activity, delaying PPARG commitment. As YAP/TAZ activity declines, cells spend less time at high ERK activity, allowing PPARG to rise and differentiation to proceed (Fig. 7). In this way, YAP/TAZ–ERK dynamics determine how many progenitor divisions occur before commitment and therefore how many differentiated cells are ultimately produced.

**Figure 7.**
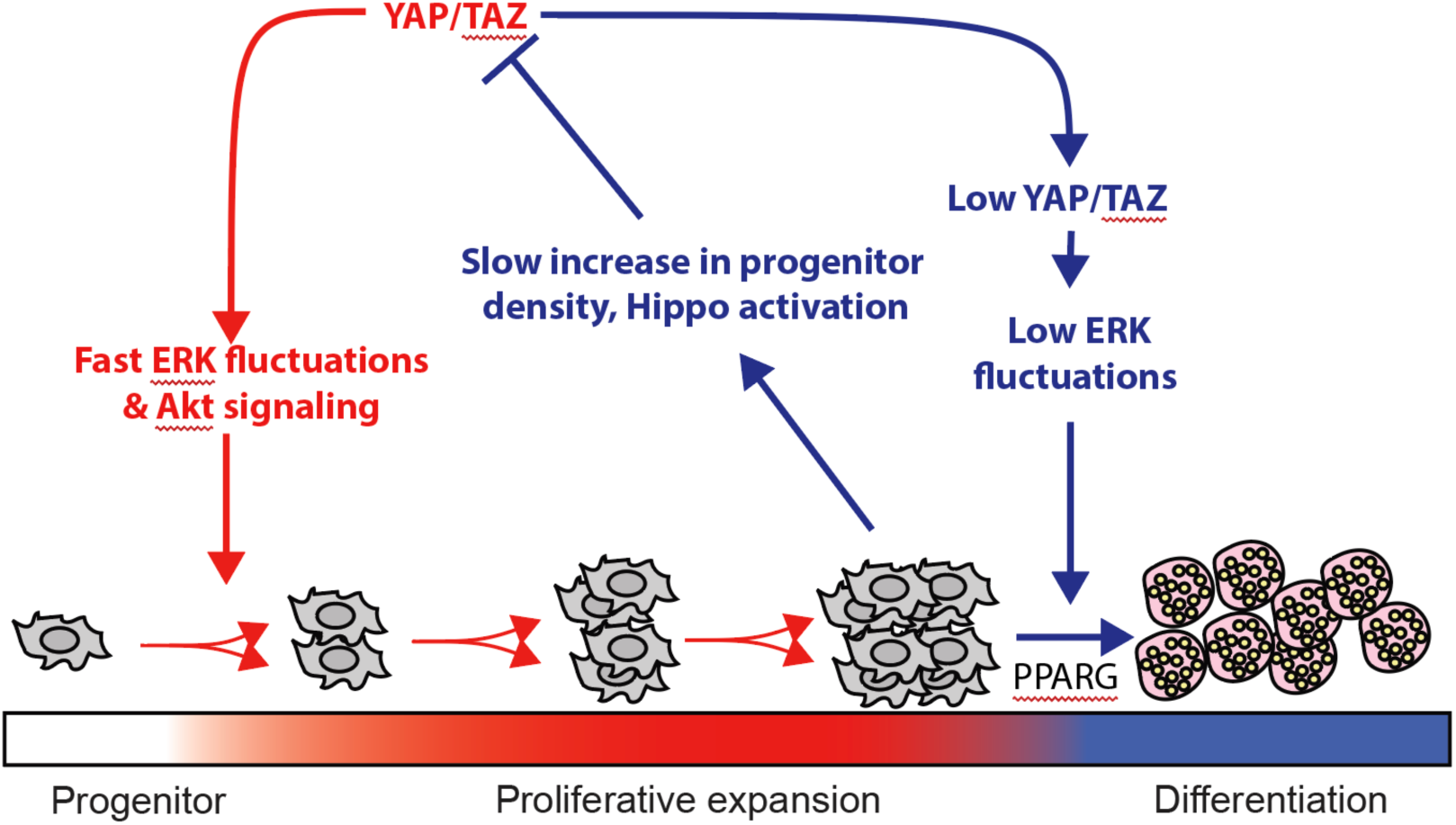
YAP/TAZ regulate ERK dynamics to coordinate progenitor expansion and differentiation commitment. YAP/TAZ promote AKT- and ERK-dependent progenitor proliferation while maintaining cells in a high-ERK state that delays PPARG accumulation. As the progenitor population expands, we propose that increasing cell density and contact-dependent Hippo/LATS signaling progressively reduce YAP/TAZ activity, providing a slow negative-feedback mechanism that terminates the proliferative phase. Declining YAP/TAZ activity causes cells to spend less time at high ERK activity, allowing PPARG to accumulate and cells to commit to differentiation. Thus, YAP/TAZ couple expansion of the progenitor pool to the subsequent transition into differentiation through fast regulation of ERK signaling and slower density-dependent negative feedback.

## Discussion

Using live single-cell imaging of PPARG, cell-cycle, and ERK-activity reporters, we identify a YAP/TAZ–ERK mechanism that dynamically couples progenitor expansion to differentiation commitment. YAP and TAZ act together to promote proliferation while increasing the fraction of time cells spend in a high-ERK state that suppresses PPARG accumulation. When YAP/TAZ activity decreases, cells spend less time at high ERK activity and PPARG rises. A transient YAP/TAZ pulse can therefore expand the progenitor pool without preventing later differentiation, increasing the number of adipocytes ultimately produced. These findings suggest a self-limiting mechanism in which YAP/TAZ-driven progenitor expansion increases cell density and contact inhibition, activating Hippo signaling, reducing YAP/TAZ activity, and permitting commitment.

YAP and TAZ act together during progenitor expansion and differentiation commitment. Only simultaneous knockdown of YAP and TAZ accelerated differentiation, consistent with functional redundancy between the two paralogs. Their transcriptional output declined during adipogenesis through two parallel mechanisms: reduced nuclear YAP/TAZ and reduced TEAD expression. At matched time points, PPARG-high cells had lower nuclear-to-cytosolic YAP and TAZ ratios than PPARG-low cells, linking reduced YAP/TAZ activity directly to differentiation state at the single-cell level. However, YAP and TAZ are not completely interchangeable. Continuous TAZ activation suppressed differentiation more strongly than YAP activation, despite similar effects on progenitor proliferation, consistent with the reported ability of TAZ to directly inhibit PPARG^17^. Thus, YAP and TAZ share a redundant role in maintaining the proliferative, high-ERK progenitor state, while TAZ may exert an additional PPARG-directed brake on differentiation. This distinction may help reconcile previous studies linking YAP/TAZ activity both to adipose tissue expansion and to inhibition of adipocyte differentiation.

ERK, rather than proliferation alone, links YAP/TAZ activity to differentiation commitment. CDK4/6 or AKT inhibition suppressed YAP/TAZ-driven proliferation without restoring differentiation, whereas MEK inhibition restored differentiation even when YAP/TAZ activity remained high. These results place YAP/TAZ upstream of the ERK inactivation step that we previously showed is required for commitment^6^. Our data further show that ERK activity fluctuates dynamically and that YAP/TAZ controls commitment by regulating the fraction of time cells spend at high ERK activity. When YAP/TAZ activity remains high, cells spend more time in the high-ERK state and PPARG accumulation is delayed. When YAP/TAZ activity falls, cells spend less time at high ERK activity, allowing PPARG to rise and commitment to proceed. Additional transcriptional or epigenetic mechanisms may also contribute to YAP/TAZ-mediated suppression of PPARG^17^.

YAP/TAZ–TEAD and ERK can influence each other through multiple mechanisms. ERK can activate TAZ in some mesenchymal cells^23^, whereas our perturbation experiments place YAP/TAZ upstream of ERK in this system. Both relationships could therefore operate in the same cells. One possible link is growth-factor receptor signaling, because YAP can increase signaling through EGFR and related receptors that activate MEK–ERK^19^. Importantly, YAP/TAZ activity is also mechanically suppressed by increasing cell density and contact inhibition through Hippo/LATS signaling^11^. This provides a potential negative-feedback mechanism for terminating the proliferative phase: YAP/TAZ promote progenitor expansion, the resulting increase in progenitor density could activate Hippo/LATS signaling, and YAP/TAZ activity would then be progressively reduced. This in turn would reduce the time cells spend at high ERK activity, allowing PPARG to rise and differentiation commitment to occur. In this model, rapid YAP/TAZ control of ERK and AKT drives progenitor expansion, whereas slower density-dependent Hippo feedback provides a mechanism for terminating that expansion and enabling commitment.

The pulse experiment demonstrates the functional consequence of this temporal control. Continuous YAP/TAZ activity expands progenitors but suppresses differentiation, whereas a transient early pulse expands the progenitor pool while still allowing later commitment. The density-dependent feedback proposed in Fig. 7 provides a mechanism by which this transient YAP/TAZ activity could arise naturally: proliferation increases progenitor density, which could progressively activate Hippo/LATS signaling and reduce YAP/TAZ activity. Thus, the duration of YAP/TAZ activity determines whether progenitor expansion is ultimately followed by terminal differentiation.

We identify ERK dynamics as the mechanism linking this temporal control to commitment. Sustained YAP/TAZ activity prolongs the fraction of time cells spend in the high-ERK state, whereas declining YAP/TAZ activity reduces high-ERK-state occupancy, allowing PPARG to rise. By controlling how many divisions occur before commitment, this mechanism determines how many differentiated cells are ultimately produced. A similar need to coordinate progenitor expansion with differentiation exists in other renewing tissues, including the intestinal epithelium, where ERK waves regulate stem-cell compartment size and epithelial patterning^24,25^. Our findings identify YAP/TAZ control of ERK dynamics as a mechanism that may broadly coordinate progenitor expansion with differentiation to regulate tissue size and regeneration.

## Methods

### Cell lines and reporter constructs

All experiments used OP9 mouse preadipocytes carrying an endogenously tagged mCitrine-PPARG2 allele, generated and characterized previously^3,10^. Nuclear marker (H2B-mTurquoise2) and ERK kinase-translocation reporter (ERK-KTR-mRuby3)^26^ constructs were introduced by lentiviral transduction using a third-generation packaging system, and reporter-positive cells were enriched by FACS or by antibiotic selection after infection.

### Cell culture

OP9 cells were cultured as previously described^3,10^ in phenol red-containing MEM-α (Thermo Fisher Scientific) supplemented with 20% FBS, 100 U/mL penicillin, 100 µg/mL streptomycin, and 292 µg/mL L-glutamine, and passaged every two days.

### Adipogenesis protocol

Cells were trypsinized and plated in 96-well optical plates (Cellvis P96-1.5H-N) at 20,000 cells/cm² in MEM-α without phenol red (R&D Systems M34750) with 10% FBS. The next day, medium was replaced with the same basal medium containing 10% FBS, 1 µM dexamethasone, 125 µM IBMX, and 1.75 nM insulin (DMI; all from Sigma-Aldrich) for 48 h, after which the DMI-containing medium was aspirated and replaced with the same medium containing 1.75 nM insulin only for a further 48 h. Differentiation was carried out in phenol red-free medium for all live-imaging experiments.

### Inducible, degradable YAP and TAZ constructs

Mouse YAP1 and TAZ coding sequences (UniProt P46938 and Q91WF0) were codon-optimized (IDT) to remove homology with endogenous transcripts and all siRNA target sites. Constitutively active mutants were generated by mutating the LATS1/2 consensus phosphorylation sites (YAP S112A/S382A and TAZ S89A/S306A; YAP-CA-S2A and TAZ-CA-S2A)^11,12^; a more strongly active YAP variant (YAP-CA-S5A) carried alanine substitutions at all five consensus sites^11^. Each construct was expressed from a doxycycline-responsive promoter as a C-terminal fusion to mRuby3 followed by a minimal auxin-inducible degron (mAID), and stably introduced into OP9 reporter cells with selection for stable integrants. Auxin-dependent degradation required co-expression of the plant F-box receptor OsTIR1, introduced as a separate constitutive construct^13^. Expression was induced with ~1 µg/mL doxycycline (lower titrated doses for the Fig. 4 pulse experiments), and degradation was triggered by adding indole-3-acetic acid (IAA) to 200 µM; loss of mRuby3 signal confirmed rapid degradation.

### siRNA-mediated gene silencing

siRNAs targeting Yap1, Wwtr1 (TAZ), Lats1, Amotl2, Nf2, and Tead1-4, together with a non-targeting control, were obtained from Qiagen and IDT (sequences in Supplementary Table 2). Cells were reverse-transfected with Lipofectamine RNAiMAX (Invitrogen): 19 µL OptiMEM was mixed with 0.5 µL of a 10 µM siRNA stock per target gene (5 pmol) and 0.5 µL RNAiMAX, incubated for 10 min at room temperature, and combined with 80 µL of culture medium containing the desired number of cells; the entire ~100 µL volume was plated into one well of a 96-well plate. The transfection mixture was left on the cells for 6 h, then aspirated and replaced with 100 µL fresh culture medium. DMI was added 24 h after transfection.

### Small-molecule inhibitors

Final concentrations were: LATS1/2 inhibitor TDI-011536 (0.15-5 µM for dose-response experiments and 200 nM for combination experiments, as indicated), pan-TEAD inhibitor GNE-7883 (2.5 µM), CDK4/6 inhibitor palbociclib (PD-0332991; 2.5 µM), MEK inhibitor PD-0325901 (200 nM), and AKT inhibitor miransertib (ARQ 092; 200 nM). For drug additions during adipogenesis, 40 µL of serum-free MEM-α containing a 5× concentration of the inhibitor was added to individual wells containing cells growing in 160 µL medium, at the time of DMI addition unless otherwise indicated. Equal volumes of DMSO served as vehicle control. Thereafter, medium containing insulin plus inhibitor or vehicle was used to replace the DMI-containing medium for the remainder of the protocol.

### Immunofluorescence and EdU

Cells were fixed with 4% PFA in PBS for 15 min at room temperature and washed three times with PBS using an automated plate washer (BioTek). Cells were permeabilized with ice-cold methanol for 10 min, washed three times with PBS, and blocked for 1 h in PBS containing 5% FBS and 0.3% Triton X-100. Primary antibodies were diluted in PBS containing 1% BSA and 0.3% Triton X-100 and incubated overnight at 4 °C: YAP (Cell Signaling Technology, D8H1X, cat. 14074, 1:1000), TAZ (Cell Signaling Technology, E8E9G, cat. 83669, 1:1000), PPARG (Santa Cruz Biotechnology, E-8, sc-7273, 1:1000), Pan-TEAD (Cell Signaling Technology, D3F7L, 1:1000), phospho-p44/42 MAPK (Thr202/Tyr204) (Cell Signaling Technology, 4370, 1:400), and total p44/42 MAPK (Cell Signaling Technology, 4696, 1:400). Cells were washed three times with PBS, incubated for 1.5 h with Alexa Fluor-conjugated anti-rabbit or anti-mouse secondary antibodies (Thermo Fisher Scientific, 1:1000) in PBS containing 1% BSA and 0.3% Triton X-100, then counterstained with DAPI for 30 min at room temperature and washed three times with PBS before imaging. For proliferation, EdU (Cayman Chemical, 20518) was added to 50 µM for 15 min before fixation and detected by copper-catalyzed click chemistry with a picolyl-azide fluorophore^27^; cells were scored EdU-positive above a threshold set from the bimodal EdU distribution. At least 5000 cells were analyzed per condition for fixed experiments.

### Microscopy

Cells were plated 24 h before imaging, and full growth medium was replaced with fresh phenol red-free MEM-α supplemented with 10% FBS and, where indicated, the adipogenic stimulus, immediately before acquisition. Because OP9 cells are migratory and reach high densities during differentiation, the plating suspension was a mosaic of cells that did or did not stably express fluorescently tagged H2B, which allows the same cells to be tracked reliably over several days; the ratio of tagged to untagged cells was 1:1 to 3:1 when a perturbation was expected to reduce proliferation and 1:3 when it was expected to increase proliferation. Imaging was performed on a Nikon Eclipse Ti2 inverted microscope with a 10× Plan Apo 0.45 NA objective and 2×2 camera binning, on a stage-top humidified incubator at 37 °C and 5% CO2. Selected fixed-cell images were acquired with a 20× Plan Apo 0.75 NA objective. Live cell-cycle experiments were acquired every 12 min, whereas ERK-KTR experiments were acquired every 6 min, in three or four fluorescence channels (CFP, YFP, RFP, iRFP) as required. Total light exposure was kept below 800 ms per time point, and four to nine non-overlapping sites were imaged per well. Nuclei were labeled with DAPI (fixed) or the H2B-mTurquoise2 reporter (live).

### Western blotting

OP9 cells were plated on 60-mm dishes at ~20,000 cells/cm² and stimulated with DMI 48 h after plating. At each time point, cells were washed twice with ice-cold PBS, scraped, and lysed in ice-cold RIPA buffer (Millipore Sigma 20-188) containing protease and phosphatase inhibitors (Thermo Fisher Scientific 78444). Lysates were held on ice for 20 min and cleared at 13,000×g for 10 min; total protein was quantified by Bradford assay, and 35-40 µg was separated per lane by SDS-PAGE. Blots were blocked in TBS containing 0.1% Tween-20 (TBST) and 10% dry milk and incubated overnight with primary antibodies diluted in TBST with 5% BSA: phospho-YAP (Ser127) (Cell Signaling Technology, cat. 13008, 1:500), total YAP (Cell Signaling Technology, D8H1X, cat. 14074, 1:1000), phospho-ERK1/2 (Thr202/Tyr204) (Cell Signaling Technology, 4370, 1:2000), total ERK1/2 (Cell Signaling Technology, 4696, 1:2000), PPARG (Cell Signaling Technology, 2443, 1:1000), and HRP-conjugated β-actin (Santa Cruz Biotechnology, sc-47778, 1:4000). Blots were washed three times in TBST over 30 min and incubated for 1.5 h at room temperature in the dark with Alexa Fluor goat anti-rabbit 680 (Thermo Fisher Scientific, 1:10,000), IRDye 800CW goat anti-mouse (LI-COR Biosciences, 1:10,000), or HRP-conjugated goat anti-mouse (Cell Signaling Technology, 1:2000) diluted in TBST with 5% BSA, washed three times over 20 min, and imaged on a LI-COR Odyssey fluorescence and chemiluminescence imager.

### Quantitative RT-PCR and RNA sequencing

Total RNA was isolated with the RNeasy Mini Kit (QIAGEN, cat. 74104), and RNA quantity and quality were assessed with a NanoDrop One spectrophotometer (Thermo Scientific, ND-ONE-W). For qPCR, 1 µg RNA was used for cDNA synthesis with the qScript cDNA Synthesis Kit (Quantabio, cat. 95047-100), followed by purification with the QIAquick PCR Purification Kit (QIAGEN, cat. 28104). qPCR was performed with BlasTaq 2× qPCR MasterMix (Applied Biological Materials, cat. G891) on a Bio-Rad OPUS 384 instrument. Each 10-µL reaction contained 5 µL 2× master mix, 1 µL of 1:10 diluted cDNA, and 4 µL of premixed 5 µM primers. Primers were designed using IDT PrimerQuest with a target amplicon length of ~250 bp and target primer melting temperature of 60-65 °C; sequences are provided in Supplementary Table 3. Fold changes were computed by the ΔCt method using Gapdh as the reference gene (mean of three biological replicates, each with three technical replicates). For RNA sequencing, strand-specific libraries were prepared from polyadenylated mRNA at seven time points (days 0-6, three biological replicates), sequenced on Illumina NextSeq, pseudo-aligned with kallisto^28^ to the mouse transcriptome (GRCm38), and analyzed as transcripts per million (TPM).

### Image processing and single-cell analysis

Image processing and analysis were performed in MATLAB R2022a (MathWorks). Automated image segmentation, single-cell tracking, and fluorescence quantification used the MACKtrack package described previously^3,6^. In fixed samples, PPARG level was quantified as the median fluorescence within the nucleus of each cell, and cells were scored PPARG-high if this value exceeded a cut-off determined from the bimodal PPARG distribution at the end of the experiment. For live imaging, the CFP channel capturing H2B-mTurquoise2 was used for nuclear segmentation and tracking. Single-cell traces were filtered to remove incomplete or mis-tracked cells according to the following criteria: cells or their daughters absent at any point during the time lapse, cells showing a large step increase or decrease in PPARG intensity relative to the preceding time point, and cells with many large fluctuations in H2B signal.

The number of divisions per cell was counted from the integrated H2B signal, which doubles across the cycle and halves at each mitosis. ERK activity was measured from the ERK-KTR-mRuby3 reporter as the ratio of cytosolic to nuclear median fluorescence. Nuclear fluorescence was measured directly within the H2B segmentation mask; cytosolic fluorescence was measured within an annulus generated by isometrically expanding the nuclear mask by 4 pixels and subtracting the nuclear mask. Cells were scored as high-ERK when this ratio exceeded a fixed threshold applied identically across all conditions.

To identify the differentiation commitment point, a PPARG cut-off was determined by scanning the entire PPARG time series for the value that maximized the number of cells satisfying all three of the following: PPARG below threshold at the start of the experiment, PPARG above threshold immediately before removal of the adipogenic stimulus, and PPARG above threshold at the end of the experiment. For each cell that remained above this threshold at the end of the experiment, the commitment time was defined as the first time PPARG crossed it.

### Statistics

Each experiment included at least two replicate wells per treatment condition, and experiments were repeated on separate days as biological replicates, as indicated in the figure legends. Live time-course data are shown as mean ± 95% confidence interval; box and violin plots show the 25th, 50th, and 75th percentiles together with the data distribution. P values are from unpaired t-tests unless otherwise noted; qPCR comparisons used two-way ANOVA. Results are representative of at least two independent experiments.

## Acknowledgements

We thank all members of the Teruel and Meyer laboratories for helpful discussions. This work was supported by National Institutes of Health grant R01-DK131432-01A1 (M.N.T.) and startup funds from the Drukier Institute for Children’s Health and Weill Cornell Medicine (M.N.T.).

## Author Contributions

S.S. and M.N.T. designed the study. S.S., H.E.B., and Z.Z. performed experiments and collected and analyzed data. T.M. gave conceptual advice. M.N.T. provided supervision and funding acquisition. S.S. and M.N.T. wrote the manuscript with input from all authors.

