## Supplementary Material for "YAP/TAZ-controlled ERK dynamics coordinate progenitor expansion and differentiation commitment"

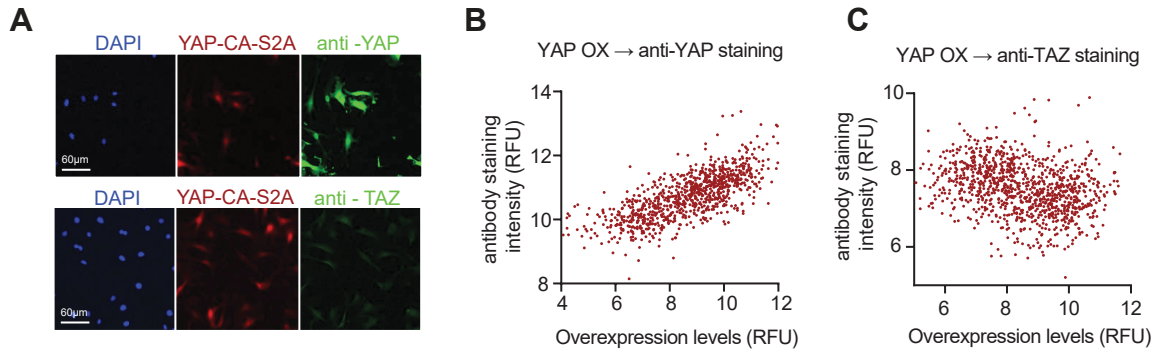

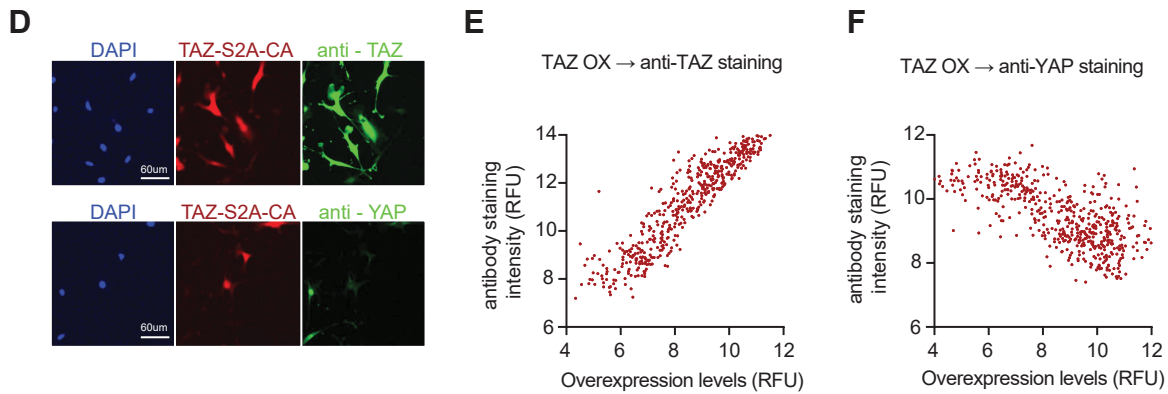

### Supplementary Fig. 1| YAP/TAZ levels are regulated through a mutual negative feedback

(a), (d) Representative immunofluorescence images of OP9 cells expressing constitutively active YAP (YAP-CA-S2A) and constitutively active TAZ (TAZ-CA-S2A) constructs respectively. Channels shown (left to right): Nuclear stain Hoechst (blue), YAP or TAZ -CA-S2A-mRuby3 (red) and antibody staining for YAP or TAZ (green). Scale bar, 60 μm.

(b), (f) Single-cell Immunofluorescence staining intensity for YAP as a function of YAP or TAZ -CA-S2A::mRuby3 levels in response to doxycycline-induced overexpression (1 μg/mL). The negative trend in (c) shows that increasing YAP-CA-S2A expression suppresses endogenous TAZ.

(c), (e) Immunofluorescence staining intensity for TAZ as a function of YAP or TAZ -CA-S2A::mRuby3 levels in response to doxycycline-induced overexpression (1 μg/mL). The negative trend in (f) shows that increasing TAZ-CA-S2A expression suppresses endogenous YAP.

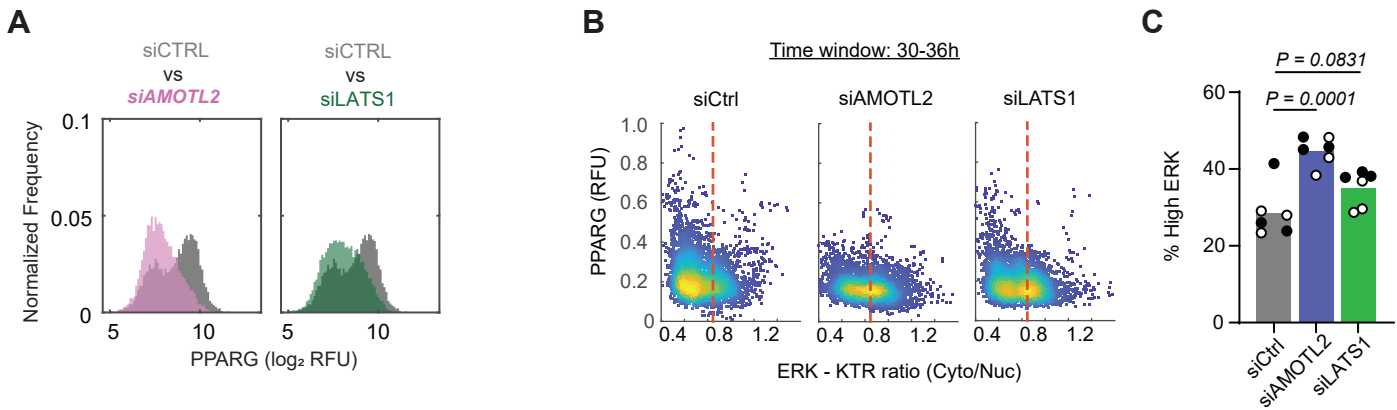

**Supplementary Fig. 2| Increasing YAP/TAZ activity increases the time cells spend high ERK activity state and delays their differentiation commitment**

**(a)** Normalized frequency histograms showing the distribution of PPARG levels (log<sub>2</sub> RFU) at 48h after DMI stimulation in AMOTL2 siRNA treated cells (siAMOTL2, pink) compared to control cells (siCTRL, gray) and in LATS1 siRNA (siLATS1, green) treated cells compared to control cells (gray). Cell numbers: siCTRL n = 7,524; siAMOTL2 n = 8,801; siLATS1 n = 10,849. Representative of 2 independent experiments.

**(b)** Single-cell scatter plots of PPARG versus ERK-KTR ratio scatter plots in the 30-36h time window since the start of DMI stimulation for siCtrl, siAMOTL2 and siLATS1 conditions. Representative of 2 independent experiments.

**(c)** Fraction of high-ERK timepoints per cell in the 30-36h. Bars show the means and individual points show the means of 6 replicate wells pooled from 2 independent experiments containing 3 wells each. The points from replicate experiments are shown as filled and open circles for each condition respectively. One way ANOVA with Dunnett's multiple comparison test. All pairwise comparisons are against the control condition.
